# Growth-independent cross-feeding modifies boundaries for coexistence in a bacterial mutualism

**DOI:** 10.1101/083386

**Authors:** Alexandra L. McCully, Breah LaSarre, James B. McKinlay

## Abstract

Nutrient cross-feeding can stabilize microbial mutualisms, including those important for carbon cycling in nutrient-limited anaerobic environments. It remains poorly understood how nutrient limitation within natural environments impacts mutualist growth, cross-feeding levels, and ultimately mutualism dynamics. We examined the effects of nutrient limitation within a mutualism using theoretical and experimental approaches with a synthetic anaerobic coculture pairing fermentative *Escherichia coli* and phototrophic *Rhodopseudomonas palustris*. In this coculture, *E. coli* and *R. palustris* resemble an anaerobic food web by cross-feeding essential carbon (organic acids) and nitrogen (ammonium), respectively. Organic acid cross-feeding stemming from *E. coli* fermentation can continue in a growth-independent manner during nutrient limitation, while ammonium cross-feeding by *R. palustris* is growth-dependent. When ammonium cross-feeding was limited, coculture trends changed yet coexistence persisted under both homogenous and heterogenous conditions. Theoretical modeling indicated that growth-independent fermentation was crucial to sustain cooperative growth under conditions of low nutrient exchange. We also show that growth-independent fermentation sets the upper *E. coli* cell density at which this mutualism is supported. Thus, growth-independent fermentation can conditionally stabilize or destabilize a mutualism, indicating the potential importance of growth-independent metabolism for nutrient-limited mutualistic communities.

**Conflict of interest:** The authors declare no conflict of interest.

## Introduction

Mutualistic cross-feeding interactions between microbes crucially impact diverse processes ranging from human health (Hammer *et al.*, 2014; Ramsey and Whiteley, 2009; Flint *et al.*, 2007) to biogeochemical cycles (Morris *et al.*, 2013; McInerney *et al.*, 2010; Durham *et al.*, 2015). Within most environments, microbial communities experience prolonged periods of nutrient limitation (Lever *et al.*, 2015). In general, bacteria tolerate nutrient limitation by modulating their growth and metabolism (Ferenci, 2001; Russell and Cook, 1995; Wanner and Egli, 1990; Rittershaus *et al.*, 2013; Lee *et al.*, 1976). Sub-optimally growing and even non-growing cells can survive by retaining low metabolic activity to generate maintenance energy (Wanner and Egli, 1990; Russell and Cook, 1995; Rittershaus *et al.*, 2013; Hoehler and Jørgensen, 2013). It is possible that growth-independent metabolic products could serve to cross-feed other microbes and thereby influence the initiation and/or endurance of microbial mutualisms under growth-limiting conditions. Nonetheless, most microbial cross-feeding studies view nutrient release as being tightly coupled to growth. While mutualism flux balance models tend to include growth-independent maintenance energy parameters (Harcombe *et al.*, 2014; Chubiz *et al.*, 2015), most other mutualism models do not, and few studies have examined the impact of growth-independent cross-feeding on mutualism dynamics (Shou *et al.*, 2007; Megee III *et al.*, 1972; Stolyar *et al.*, 2007).

Growth-independent metabolism is fundamental to fermentative microbes during nutrient limitation, as they must continue to ferment and excrete products, albeit at a lower rate than during growth, to generate maintenance energy. For example, in the absence of electron acceptors, and starved for essential elements (i.e., nitrogen or sulfur), *Escherichia coli* generates energy by fermenting glucose in a growth-independent manner (Wanner and Egli, 1990; LaSarre *et al.*, 2016). Fermentative microbes serve pivotal roles within natural anaerobic food webs, wherein their excreted products serve as nutrients for other microbes. As such, growth-independent fermentation could be an important cross-feeding mechanism within anaerobic communities.

Studying mutualistic cross-feeding in natural environments can be challenging due to environmental and genetic stochasticity. Synthetic microbial communities, or cocultures, offer an alternative approach that mimics key aspects of natural communities while providing a greater degree of experimental control (Widder *et al.*, 2016; Momeni *et al.*, 2011; Ponomarova and Patil, 2015; Lindemann *et al.*, 2016). We previously developed a bacterial coculture to facilitate the study of mutualistic cross-feeding in anaerobic environments (LaSarre *et al.*, 2016). Our coculture resembles other fermenter-photoheterotroph cocultures, which have primarily been studied for converting plant-derived sugars into H_2_ biofuel (Odom and Wall, 1983; Fang *et al.*, 2006; Sun *et al.*, 2010; Ding *et al.*, 2009). However, unlike previous systems, our coculture enforces stable coexistence through bi-directional cross-feeding of essential nutrients. Specifically, *E. coli* ferments sugars to excreted organic acids, providing essential carbon and electrons for a genetically engineered *Rhodopseudomonas palustris* strain (Nx). *R. palustris* Nx has a NifA* mutation (McKinlay and Harwood, 2010a) that results in NH_4_^+^ excretion during N_2_ fixation, providing essential nitrogen for *E. coli* (Figure 1) (LaSarre *et al.*, 2016). We previously used our coculture to examine the effects of increased NH_4_^+^ cross-feeding on coculture dynamics (LaSarre *et al.*, 2016). In that study, theoretical modeling suggested that coexistence would persist even at very low NH_4_^+^ excretion levels (LaSarre *et al.*, 2016). This prediction prompted us to ask herein: how does this mutualism contend with limitation of cross-fed nutrients?

**Figure 1.**
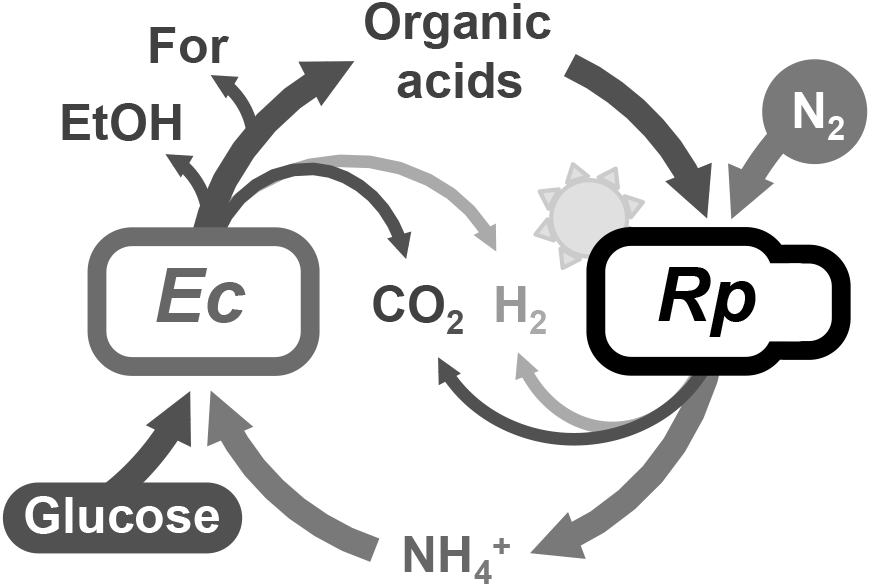
**Bi-directional anaerobic cross-feeding between fermentative *Escherichia coli* and phototrophic *Rhodopseudomonas palustris* Nx.** Obligate bi-directional cross-feeding of carbon to *R. palustris* (*Rp*) and nitrogen to *E. coli* (*Ec*) imparts stable coexistence. *E. coli* anaerobically ferments glucose into fermentation products, including the organic acids, acetate, lactate, and succinate, which provide essential carbon for *R. palustris* Nx. In return *R. palustris* Nx uses light energy to fix N_2_ and excrete NH_4_^+^, which provides *E. coli* with essential nitrogen. Formate and ethanol, produced by *E. coli*, and CO_2_ and H_2_, produced by both *E. coli* and *R. palustris*, accumulate in cocultures. *E. coli* fermentation can be growth-independent. Filled bubbles indicate compounds added exogenously to the coculture.

Using theoretical and experimental approaches, we show that growth-independent fermentation is crucial for maintaining cooperative growth during limitation of cross-fed NH_4_^+^. Conversely, we also show that growth-independent fermentation can be detrimental to the mutualism under certain circumstances. Specifically, growth-independent fermentation can inhibit coculture growth when large *E. coli* populations magnify the otherwise low growth-independent fermentation rate; this leads to organic acid production that outpaces consumption and ultimately acidifies the environment and halts growth. Thus, growth-independent cross-feeding conditionally influences this mutualism in positive and negative manners and thereby sets both the upper and lower thresholds for cooperation.

## Materials and Methods

### Strains, plasmids, and growth conditions

Strains are listed in Supplementary Table 1. *E. coli* and *R. palustris* were cultivated on Luria-Burtani (LB) agar or defined mineral (PM) (Kim and Harwood, 1991) agar with 10 mM succinate, respectively. For determining colony forming units (CFU), LB agar or PM agar minus (NH4)2SO4 were used for *E. coli* and *R. palustris*, respectively. Cultures were grown in 10-mL of defined M9-derived coculture medium (MDC) (LaSarre *et al.*, 2016) in 27-mL anaerobic test tubes. The medium was made anaerobic by bubbling with N_2_, sealed with rubber stoppers and aluminum crimps, and then autoclaved. After autoclaving, MDC was supplemented with cation solution (1 % v/v; 100 mM MgSO4 and 10 mM CaCl2) and glucose (25 mM). For defined N_2_ concentrations, the medium was bubbled with argon and after autoclaving defined volumes of N_2_ were injected through a 0.2 micron syringe filter. All cultures were grown at 30°C either laying horizontally under a 60 W incandescent bulb with shaking at 150 rpm (shaking conditions) or upright without agitation (static conditions). Static cultures were only mixed for sampling upon inoculation and at the termination of an experiment. Thus, growth rates were not measured under static conditions. Starter cultures were inoculated with 200 μL MDC containing a suspension of a single colonies of each species. Test cocultures were inoculated using a 1% inoculum from starter cocultures. For serial transfers, cocultures were incubated for either one week (shaking), two weeks (100% N_2_; static), or four weeks (18% N_2_; static) before transferring a 1% stationary phase inoculum to fresh medium.

### Analytical procedures

Cell density was assayed by optical density at 660 nm (OD_660_) using a Genesys 20 visible spectrophotometer (Thermo-Fisher, Waltham, MA, USA). Growth curve readings were taken in culture tubes without sampling. Specific growth rates were determined using measurements between 0.1-1.0 OD_660_ where there is linear correlation between cell density and OD_660_. Final OD_660_ measurements were taken in cuvettes and samples were diluted into the linear range as necessary. To compare cell densities between growth conditions, CFUs were converted into growth yields by calculating CFUs per μmol glucose consumed, as N_2_ limitation prevented complete glucose consumption during the assay period. H_2_ and N_2_ were quantified using a Shimadzu (Kyoto, Japan) gas chromatograph (GC) with a thermal conductivity detector as described (Huang *et al.*, 2010). Glucose, organic acids, and ethanol were quantified using a Shimadzu high-performance liquid chromatograph (HPLC) as described (McKinlay *et al.*, 2005). NH_4_^+^ was quantified using an indophenol colorimetric assay as described (LaSarre *et al.*, 2016).

### Mathematical modeling

A Monod model describing bi-directional cross-feeding in batch cultures, called SyFFoN_v2 (Syntrophy between Fermenter and Fixer of Nitrogen), was modified from our previous model (LaSarre *et al.*, 2016) as follows: (i) a sigmoidal function, rather than a Monod function, was used to control the transition to growth-independent fermentation (10/(10+1.09^(1000*uEc)^)); (ii) sigmoidal functions were used to transition from NH_4_^+^ excretion (1-(40/(40+1.29^N^))) to H_2_ production (40/(40+1.29^N^)) by *R. palustris* as N_2_ becomes limiting; (iii) a sigmoidal function was used to simulate the inhibiting effects of accumulated organic acids on both growth and metabolism for both species (b_*x*_/(b_*x*_+10^(f+C)^)); (iv) a sigmoidal function was used to dampen growth-independent fermentation rates when consumable organic acids (lactate, succinate, and acetate) accumulate (r_*x*_^*^(100/(100+6^C^)) + r_*x*__mono) and simulate the slow growth-independent fermentation observed in *E. coli* monocultures (LaSarre *et al.*, 2016), compared to faster growth-independent fermentation in coculture; (v) *R. palustris* H_2_ production was coupled to consumable organic acid depletion, assuming that 0.5 CO_2_ are produced per H_2_ (McKinlay *et al.*, 2014); (vi) the *R. palustris* Km for N_2_ (KN) was given a value of 6 mM, based on the change in growth rate at limiting N_2_ concentrations in coculture; (vii) product formation parameters (R and r) were increased to more accurately simulate observed growth rates in coculture; (viii) the *E. coli* acid resistance parameter (bEc) was increased relative to that for *R. palustris* (b_Rp_) based on terminal pH values observed in *E. coli* monocultures versus cocultures. Equations are listed below with default values in Supplementary Table 2. SyFFoN_v2 runs in R studio and is available for download at: https://github.com/McKinlab/Coculture-Mutualism.

Equations 1 and 2 were used to describe *E. coli* and *R. palustris* growth rates:

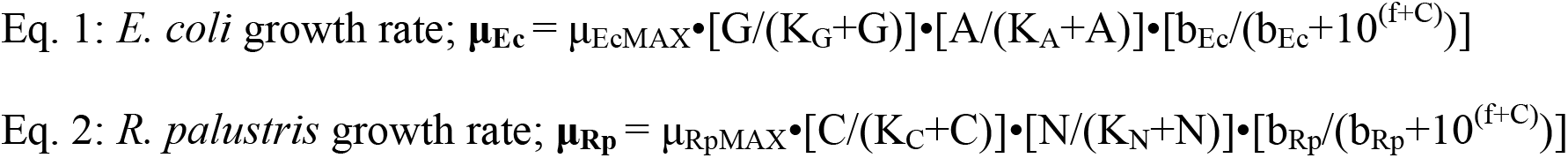

Equations 3-12 were used to describe temporal changes in cell densities and extracellular compounds. Numerical constants in product excretion equations are used to account for molar stoichiometric conversions. Numerical constants used in sigmoidal functions are based on those values that resulted in simulations resembling empirical trends. All R and r parameters are expressed in terms of glucose consumed except for RA, which is the amount of NH_4_^+^ produced per *R. palustris* cell (Supplementary Table 2).

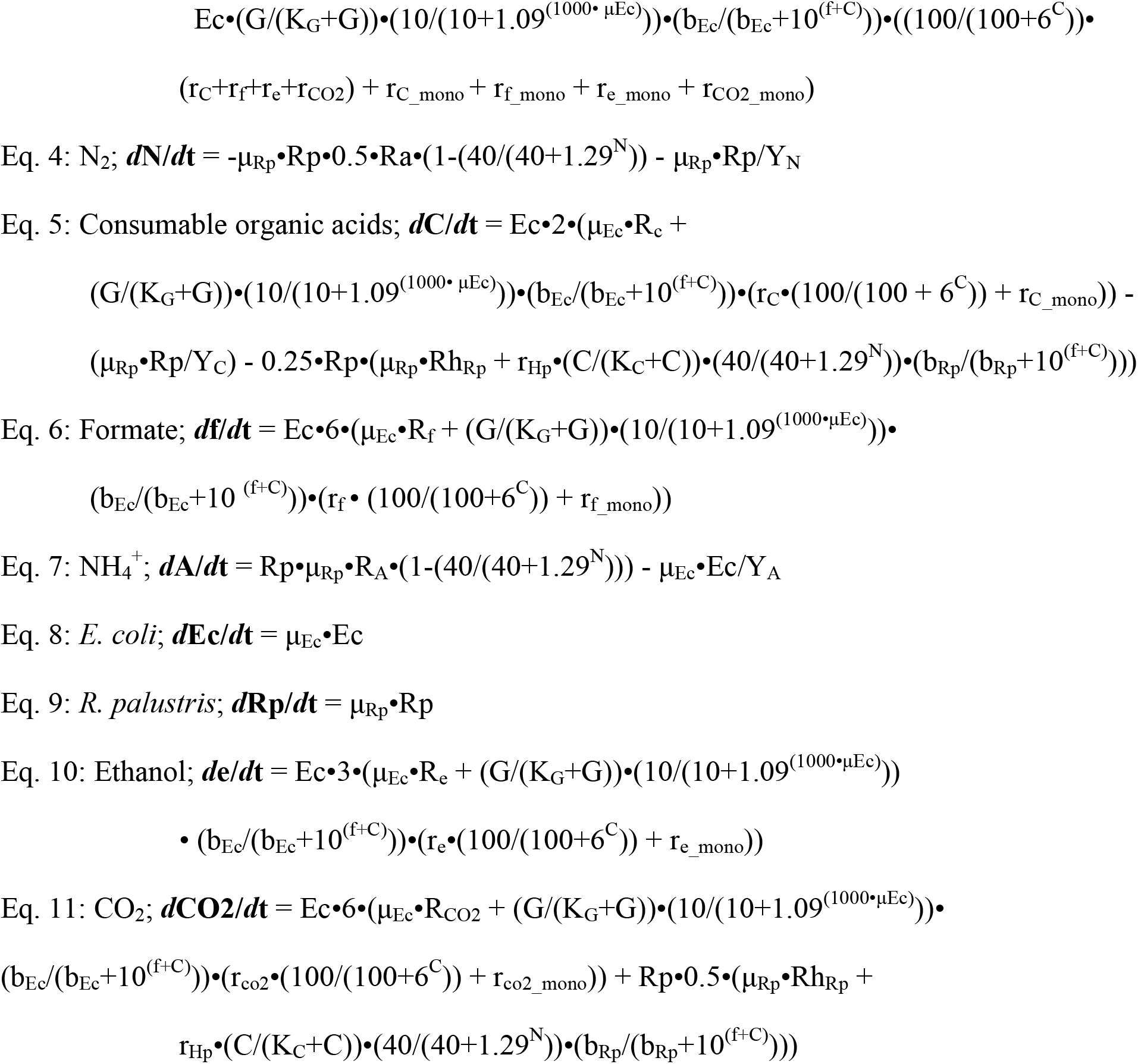
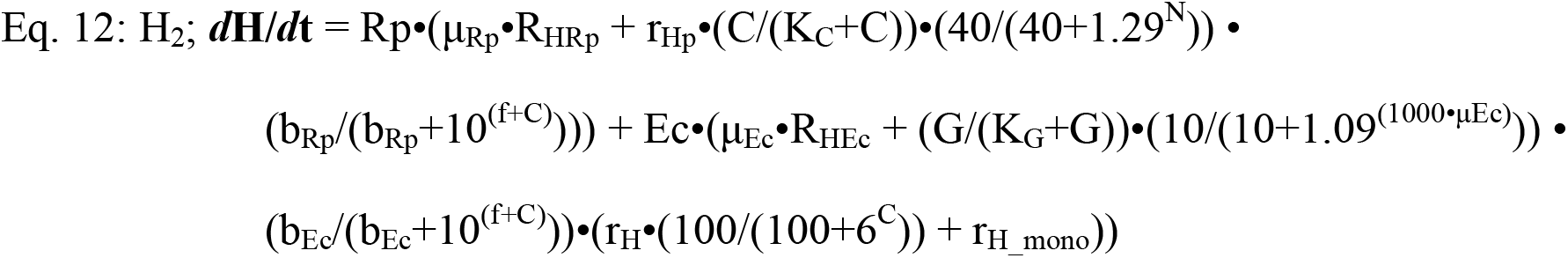

Where, μ is the specific growth rate of the indicated species (h^-1^). μMAX is the maximum specific growth rate of the indicated species (h^-1^). G, A, C, N, f, e, H and CO_2_ are the concentrations (mM) of glucose, NH_4_^+^, consumable organic acids, N_2_, formate, ethanol, H_2_, and CO_2_, respectively. All gasses are assumed to be fully dissolved. Consumable organic acids are those that *R. palustris* can consume, namely, lactate (3 carbons), acetate (2 carbons), and succinate (4 carbons). All consumable organic acids were simulated to have three carbons for convenience. Only net accumulation of formate, ethanol, CO_2_ and H_2_ are described in accordance with observed trends. K is the half saturation constant for the indicated substrate (mM). Ec and Rp are the cell densities (cells/ml) of *E. coli* and *R. palustris*, respectively. b is the ability of a species to resist the inhibiting effects of acid (mM). Y is the *E. coli* or *R. palustris* cell yield from the indicated substrate (cells / μmol glucose). Y values were determined in MDC with the indicated substrate as the limiting nutrient. R is the fraction of glucose converted into the indicated compound per *E. coli* cell during growth (μmol of glucose / *E. coli* cell), except for RA. Values were adjusted to accurately simulate product yields measured in cocultures and in MDC with and without added NH4Cl. RA is the ratio of NH_4_^+^ produced per *R. palustris* cell during growth (μmol / *R. palustris* cell). The default value was based on that which accurately simulated empirical trends. r is the growth-independent rate of glucose converted into the indicated compound (μmol / cell / h). Default values are based on those which accurately simulated empirical trends in coculture. r_mono is the growth-independent rate of glucose converted into the indicated compound by *E. coli* when consumable organic acids accumulate. Default values are based on linear regression of products accumulated over time in nitrogen-free cell suspensions of *E. coli* (LaSarre *et al.*, 2016).

## Results

### Coexistence is maintained at reduced NH_4_^+^ cross-feeding levels

Previously, we found that stable coexistence and reproducible trends in our mutualistic coculture were dependent on the transfer of NH_4_^+^ from *R. palustris* Nx to *E. coli* (LaSarre *et al.*, 2016). Adding NH_4_^+^ to the medium broke the dependency of *E. coli* on *R. palustris* and resulted in *E. coli* domination due to its higher intrinsic growth rate relative to that of *R. palustris*. Thus, the NH_4_^+^ cross-feeding level controls the *E. coli* growth rate within the mutualism. We were intrigued that theoretical modeling predicted mutualism coexistence would be maintained even at very low NH_4_^+^ cross-feeding levels, as such levels should severely limit or event halt *E. coli* growth (Supplementary Figure 1a) (LaSarre *et al.*, 2016).

To test this prediction, we sought to experimentally manipulate *R. palustris* NH_4_^+^ excretion. In our previous study, cocultures were grown under a 100% N_2_ headspace (LaSarre *et al.*, 2016). We reasoned that limiting the N_2_ supply could lower the amount of NH_4_^+^ excreted as *R. palustris* Nx would potentially retain more NH_4_^+^ for itself and excrete less for *E. coli*. To limit the N_2_ concentration, we injected known volumes of N_2_ into coculture tubes with an argon-filled headspace. In agreement with our expectation, supernatants from *R. palustris* monocultures with 18% N_2_ contained half as much NH_4_^+^ compared to 100% N_2_ monocultures (Figure 2). Thus, we concluded that N_2_ limitation was a suitable approach to manipulate NH_4_^+^ excretion levels.

To examine the degree of N_2_ limitation that would support coexistence, we grew cocultures with a range of N_2_ concentrations and monitored H_2_ yields and growth rates. We used H_2_ yield as a proxy for N_2_ limitation because H_2_ is an obligate product of nitrogenase, even under 100% N_2_ (Eq 1). As N_2_ becomes limiting, nitrogenase produces more H_2_, eventually producing pure H_2_ in the absence of N_2_ (Eq 2) (McKinlay and Harwood, 2010b; Gest and Kamen, 1949).

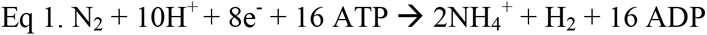

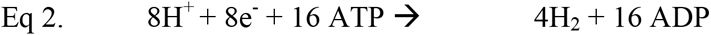

**Figure 2.**
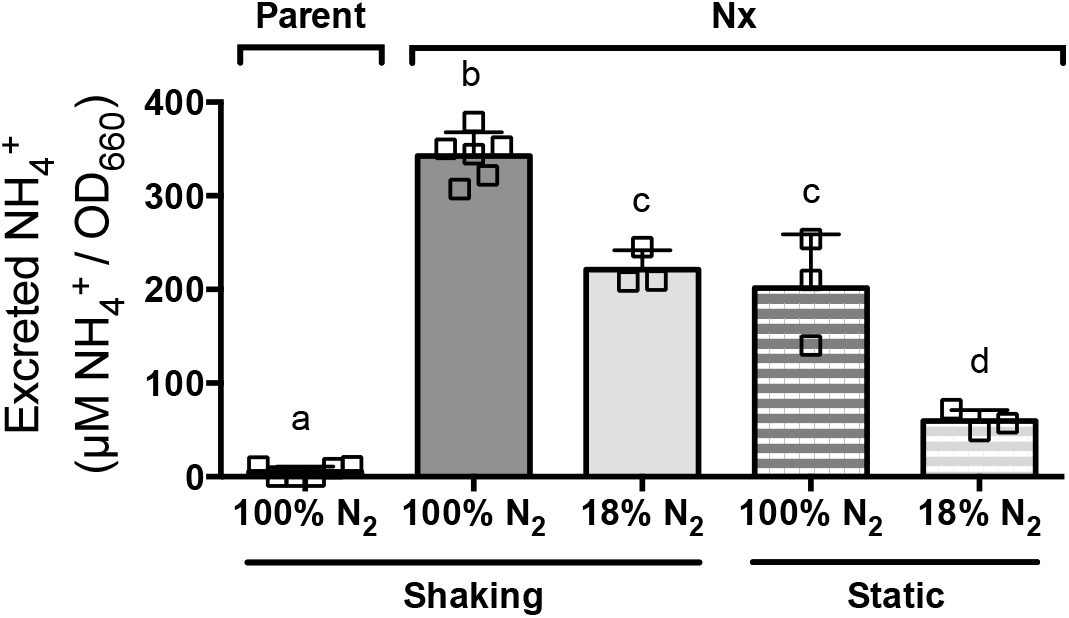
**Limiting N_2_ in monoculture results in decreased NH_4_^+^ excretion.** NH_4_^+^ in stationary-phase *R. palustris* CGA4004 (Parent) and CGA4005 (Nx) monoculture supernatants normalized to cell densities (OD_660_). *R. palustris* was cultured in MDC with 5 mM acetate either horizontally with shaking (shaking), or upright without shaking (static). Error bars indicate SD n=3-6. Different letters indicate statistical differences, p < 0.1, determined by one-way ANOVA with Tukey's multiple comparisons post test.

Thus, progressively more nitrogen limitation should result in progressively more H_2_ produced. N_2_-limited cocultures were incubated horizontally with shaking to promote gas mixing. As expected, the coculture H_2_ yield increased as N_2_ concentration decreased (Figure 3a). Nitrogen limitation was also evident from the coculture growth rate, which decreased as the N_2_ concentration decreased (Figure 3b; Supplementary Figure 2a). Notably, cocultures still grew at the lowest concentration of N_2_ that we tested, 6% N_2_ in the headspace, indicating that sufficient NH_4_^+^ was released to permit mutualistic growth.

**Figure 3.**
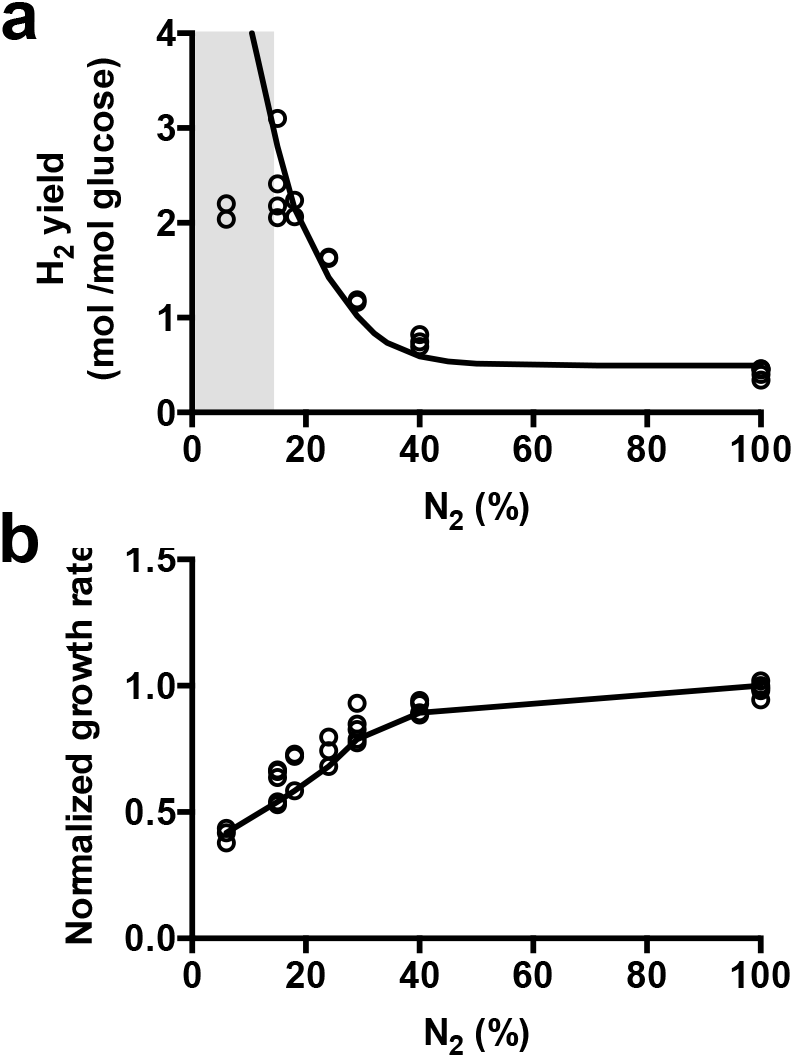
**Cocultures maintain coexistence despite reduced NH_4_^+^ exchange.** H_2_ yields (**a**) and normalized growth rates (**b**) of cocultures grown with various N_2_ concentrations (% of gas in 17 ml of headspace) under shaking conditions. Circles indicate empirical data. Lines indicate SyFFoN_v2 model predictions. (**a**) Shaded region indicates low N_2_ concentrations where empirical trends do not match model predictions. (**b**) Empirical and simulated growth rates are normalized to the corresponding average measured growth rate or simulated growth rate with 100% N_2_ (parameter N = 70 mM).

Moving forward, we focused on 18% N_2_ to characterize how lower NH_4_^+^ cross-feeding affected coculture dynamics (Figure 4). Model simulations predicted that less NH_4_ cross-feeding would result in a decrease in the *E. coli* population (Supplementary Figure 1a), In agreement with this, we observed that *E. coli* made up 5% of the population in the cocultures with 18% N_2_, which was significantly lower than the 9% *E. coli* frequency observed in cocultures with 100% N_2_. To assess coculture reproducibility, we also performed serial transfers of cocultures with 18% N_2_. Growth yield, H_2_ yield, and growth rates were all reproducible across serial transfers (Figure 4). This reproducibility indicated that coexistence was stable despite the lower level of NH_4_^+^ sharing.

**Figure 4.**
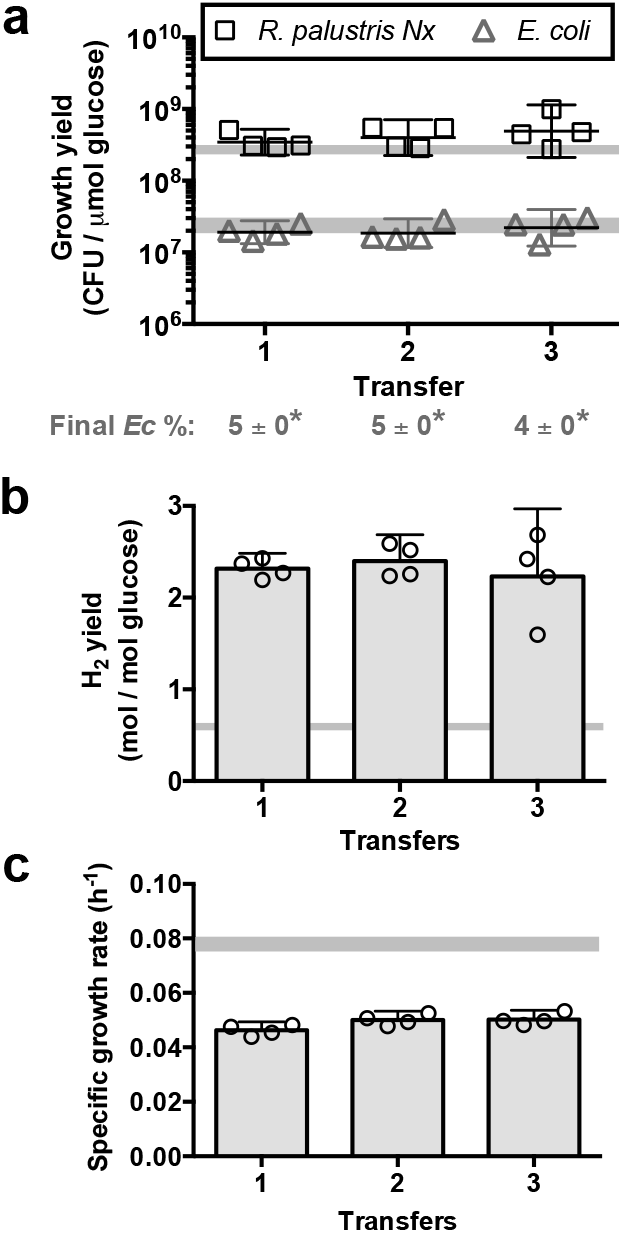
**N_2_-limited cocultures exhibit stable coexistence and reproducible trends through serial transfers in a well-mixed environment.** Individual species growth yields and final *E. coli* percentages (**a**), H_2_ yields (**b**), and growth rates (**c**) from three serial transfers of shaken cocultures grown with 18% N_2_. Transfer 1 was inoculated from a stationary phase starter coculture grown under 100% N_2_ with shaking. Cocultures were serially transferred every 7 days. Error bars indicate 95% CI, n=4. Shaded horizontal bars indicate 95% CI for shaken cocultures grown with 100% N_2_ (LaSarre *et al.*, 2016). (**a**) Growth yields are compared for each species rather than final cell densities to account for incomplete glucose consumption after 1 week in cocultures with 18% N_2_. Final *E. coli* percentages are the mean ± SD, n=4. *, statistical difference from *E. coli* percentages in shaken cocultures serially transferred with 100% N_2_ (LaSarre *et al.*, 2016), p < 0.0001, determined using one-way ANOVA with Tukey's multiple comparison post test.

### Coexistence is maintained in heterogeneous environments that decrease NH_4_^+^ cross-feeding

Spatial structuring can profoundly impact microbial mutualistic interactions. In some cases, defined spatial structuring is important or even required for coexistence (Summers *et al.*, 2010; Hom and Murray, 2014; Harcombe, 2010; Kim *et al.*, 2008). In other cases, well-mixed environments are sufficient to promote cooperative relationships (Pande *et al.*, 2014; Hillesland and Stahl, 2010; Mee *et al.*, 2014). We hypothesized that the homogeneous environment in our shaking cocultures might dampen the impact of low NH_4_^+^ cross-feeding levels. Thus, next we examined if a heterogeneous environment would affect coexistence within nitrogen-limited cocultures.

One way to induce a heterogeneous environment is by incubating in static conditions, wherein cocultures are not agitated. Static incubation was expected to result in a gradient of N_2_ availability throughout the depth of the coculture, as N_2_ has to diffuse from the headspace into the liquid. Static incubation also resulted in substantial cell settling within the coculture. Thus, we hypothesized that a heterogeneous environment would develop wherein some cells would experience a higher degree of nitrogen limitation than others, and *R. palustris* NH_4_^+^ excretion levels would thus vary. Confirming this hypothesis, static *R. palustris* Nx monocultures with 100% N_2_ showed less NH_4_^+^ excretion than in shaken cultures (Figure 2). Furthermore, in static *R. palustris* Nx monocultures with only 18% N_2_, NH_4_^+^ was only detectable after a longer 4-week incubation time (Figure 2). Thus, under static conditions, consumption of N_2_ by *R. palustris* likely exceeds the rate at which dissolved N_2_ is replenished from the headspace, resulting in N_2_ limitation and thereby low NH_4_^+^ excretion levels.

To determine how heterogeneous environments affected coculture trends under NH_4_^+^ cross-feeding limitation, we performed serial transfers of cocultures under static conditions with either 100% or 18% N_2_ in the headspace, every 2 or 4 weeks, respectively. These longer incubation times were necessary to achieve similar final cell densities between shaking and static environments. Static cocultures with 100% N_2_ had higher H_2_ yields than shaken cocultures (Figure 5a). This was expected given that a subset of the static coculture was experiencing N_2_ limitation (see Equations 1 and 2). Supplying only 18% N_2_ amplified this trend further (Figure 5a). In agreement with previously modeled effects of decreased NH_4_^+^ cross-feeding (Supplementary Figure 1a), *R. palustris* growth yields remained similar under all conditions whereas *E.* coli growth yields decreased in static cocultures relative to shaken cocultures with 100% N_2_ (Figure 5b). The decrease in *E. coli* growth yield was exacerbated in static cocultures with 18%, wherein NH_4_^+^ cross-feeding levels should be even lower (Figure 2). Coexistence was maintained over serial transfers regardless of N_2_ availability (Figure 5b). Collectively, these data demonstrate the robustness of our coculture to low NH_4_^+^ cross-feeding levels in both homogenous and heterogeneous environments.

**Figure 5.**
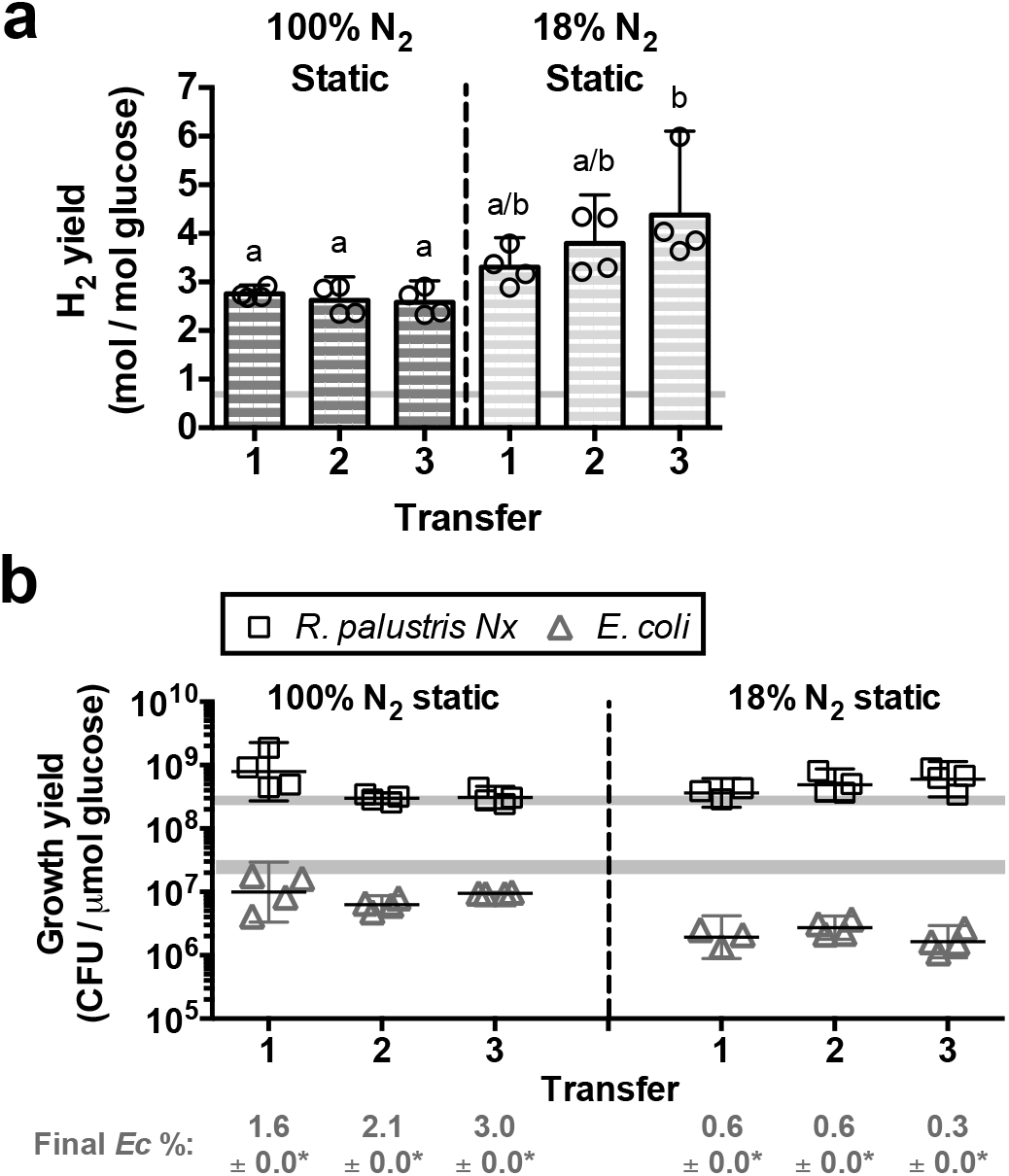
**N_2_-limited cocultures exhibit stable coexistence and reproducible trends through serial transfers in a heterogenous environment.** H_2_ yields (**a**), and individual species growth yields and final *E. coli* percentages (**b**) from three serial transfers of static cocultures grown with either 100% or 18% N_2_. Transfer 1 was inoculated from a stationary phase starter coculture grown under 100% N_2_ with shaking. Static cocultures with 100% N_2_ and 18% N_2_ were serially transferred every 2 and 4 weeks, respectively. Error bars indicate 95% CI, n=4. Shaded horizontal bars indicate 95% CI for shaken cocultures grown with 100% N_2_ (LaSarre *et al.*, 2016). (**a**) Different letters indicate statistical differences, p <0.01, determined by one-way ANOVA with Tukey's multiple comparisons post test;. (**b**) Final *E. coli* (Ec) percentages are the mean ± SD, n=4; *statistical difference from *E. coli* percentages in shaken cocultures serially transferred with 100% N_2_ (LaSarre *et al.*, 2016), p < 0.0001, determined using one-way ANOVA with Tukey's multiple comparison post test.

### Growth-independent fermentation is crucial for coexistence at low cross-feeding levels

Limiting NH_4_^+^ cross-feeding resulted in lower *E. coli* growth yields but had no effect on *R. palustris* growth yields (Figure 5). *R. palustris* growth yields depend on carbon acquisition from *E. coli*. Thus, despite lower *E. coli* cell densities during low NH_4_^+^ cross-feeding, similar levels of organic acids were still provided to *R. palustris*. We hypothesized that this disparity in growth yields could be due to growth-independent fermentation by *E. coli*. By this hypothesis, NH_4_^+^-limited *E. coli* would grow at a slower rate but would continue to use fermentation for maintenance energy; consequently, *R. palustris* would receive a slower but continuous supply of organic acids for growth and N_2_ fixation. Ultimately, *E. coli* would assimilate less glucose as a larger proportion would be used for maintenance whereas *R. palustris* would receive a similar or even greater amount of carbon from *E. coli*. This hypothesis in turn implies that growth-independent fermentation by *E. coli* is important for sustaining *R. palustris* metabolism and thereby coculture viability under low cross-feeding levels.

Growth-independent fermentation is essential for *E. coli* maintenance energy and is intimately tied to central metabolism. Thus, rather than try to genetically eliminate growth-independent fermentation we instead used a modeling approach to gauge its importance during N_2_ limitation. To simulate the impact of N_2_ limitation on coculture dynamics we first modified our previous model (LaSarre *et al.*, 2016) to account for the effects of N_2_ limitation on the shift from NH_4_^+^ to H_2_ production by *R. palustris* (SyFFoN_v2; Supplementary Table 2). We then adjusted SyFFoN_v2 parameters to simulate growth rate and metabolite yield data observed at various N_2_ concentrations (Figure 3 and Supplementary Figure 2, lines). In doing so we found that parameters based on *E. coli* monoculture data (LaSarre *et al.*, 2016) could not accurately simulate coculture growth rates observed at low N_2_ concentrations. Rather, *E. coli* growth-independent fermentation levels had to be increased by up to two-orders of magnitude to accurately simulate empirical growth rates (Figure 3b) (Supplementary Table 2). The need for these changes suggests that *R. palustris* consumption of fermentation products pulls *E. coli* fermentation by minimizing end-product inhibition, analogous to what has been observed in other fermentative cross-feeding systems (Iannotti *et al.*, 1973; Hillesland and Stahl, 2010). SyFFoN_v2 accurately predicted H_2_ yields (Figure 3a), normalized growth rates (Figure 3b), and product yields (Supplementary Figure 2) between 15% and 100 % N_2_. Below 15% N_2_, *R. palustris* likely transitions into a physiological starvation response that our model does not predict, such as the diversion of resources to storage products (McKinlay *et al.*, 2014). We also verified that SyFFoN_v2 could reproduce trends from our previous study, (LaSarre *et al.*, 2016), namely the effects of added NH_4_^+^ (Supplementary Figure 3) and varying the *R. palustris* NH_4_^+^ excretion levels (Supplementary Figure 1a).

To examine how growth-independent fermentation influenced this mutualism, we used SyFFoN_v2 to simulate the effect of N_2_ limitation on population dynamics in the presence or absence of growth-independent fermentation (Figure 6). With growth-independent fermentation included, the model predicted that mutualistic growth would be sustained at low N_2_ concentrations. *E. coli* final cell densities were predicted to decline as N_2_ levels fall below ~30% while *R. palustris* final cell densities would decline as N_2_ levels fall below ~20% (Figure 6a). Without growth-independent fermentation, simulations predicted a truncated range of N_2_ concentrations that would support coculture growth (Figure 6b). In fact, the simulations suggested that growth-independent fermentation is necessary at N_2_ concentrations where we observed reproducible coculture growth trends (Figure 6). The model predicted similar trends when NH_4_^+^ excretion levels were varied in place of N_2_ availability (Supplementary Figure 1).

**Figure 6.**
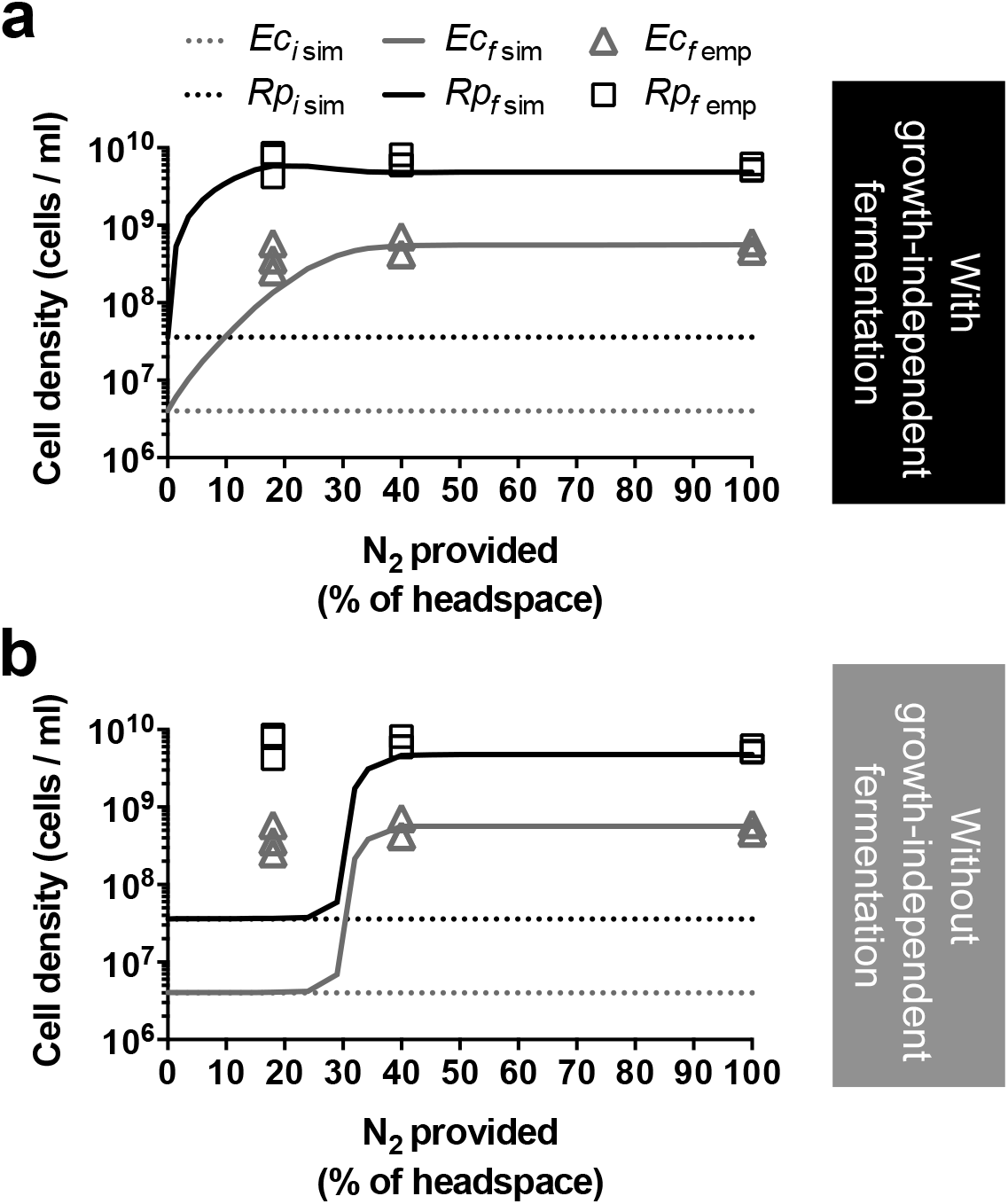
**Growth-independent fermentation permits coexistence at low NH_4_^+^ cross-feeding levels.** Simulated cell densities (lines) from cocultures grown with different N_2_ concentrations when growth-independent fermentation is included (**a**) or omitted (**b**) from the model. Ec*i* sim and Rp*i* sim, initial simulated *E. coli* (*Ec*) and *R. palustris* (*Rp*) cell densities; Ec*f* sim and Rp*f* sim, final simulated *E. coli* and *R. palustris* cell densities. Symbols are empirical CFUs/mL data for *E. coli* (Ec*f* emp) and *R. palustris* (Rp*f* emp) from 7 day samples from shaken cocultures with either 18%, 40% or 100% N_2_. The same empirical data is overlaid on each panel (n=3).

A closer inspection of simulated cross-feeding levels revealed why coculture growth would not be supported at low N_2_ concentrations (or low NH_4_^+^ cross-feeding levels) without growth-independent fermentation. At high N_2_ levels (100% N_2_), growth-coupled fermentation alone is sufficient to support coculture growth, as any increase in populations results in progressively more metabolites exchanged over time (Supplementary Figure 4a). However, near the transitional N_2_ concentration where coculture growth is predicted to fail in the absence of growth-independent fermentation (28% N_2_), metabolite excretion levels decrease as populations grow, resulting in continuously less essential resources for subsequent generations despite available glucose; in other words, cross-feeding spirals into a cycle of diminishing returns (Supplementary Figure 4b). Our data indicate that growth-independent fermentation can circumvent diminishing returns. Fermentation products will always be produced, and thus *R. palustris* will eventually grow to a density that collectively excretes sufficient NH_4_^+^ to allow *E. coli* growth. Indeed, when growth-independent fermentation is included at 28% N_2_, growth-independent cross-feeding by *E. coli* stimulates sufficient reciprocal NH_4_^+^ excretion to sustain coculture growth (Supplementary Figure 4c). These simulations strongly suggest that growth-independent fermentation permits cooperative growth at low NH_4_^+^ excretion levels that would otherwise be insufficient.

### Growth-independent metabolism prevents cooperative growth at high *E. coli* cell densities

On a per cell basis, growth-independent fermentation is considerably slower than fermentation during growth (Russell and Cook, 1995). However, we reasoned that a high *E. coli* cell density could amplify this low rate such that organic acid production would be substantial at a population level. We previously demonstrated that dose-dependent toxicity governs mutualism dynamics in our coculture; specifically, organic acids play a beneficial role as a carbon source for *R. palustris*, but a detrimental role when they accumulate enough to acidify the medium (LaSarre *et al.*, 2016). Thus, we hypothesized that if *E. coli* cell densities were sufficiently high, the collective growth-independent fermentation rate might destabilize the mutualism by producing organic acids faster than the smaller *R. palustris* population could consume them, resulting in coculture acidification and growth inhibition.

To test this hypothesis, we first simulated coculture growth from different initial species densities. The model correctly predicted that a common equilibrium would be reached from a wide range of initial *E. coli* densities (Figure 7a) (LaSarre *et al.*, 2016). However, in agreement with our hypothesis the model also predicted a maximum initial *E. coli* density that would allow cooperative growth (Figure 7a). An upper limit was experimentally verified, albeit at a lower *E. coli* density than what was predicted (Figure 7a. At an initial *E. coli* density of ~2 x 10^9^ CFU / ml, the pH reached acidic levels known to prevent *R. palustris* growth and metabolism (LaSarre *et al.*, 2016) (Figure 7a). As a result, neither species' population increased (Figure 7a). These results contradicted predictions when growth-independent fermentation was omitted from the model, as there was no predicted initial *E. coli* density that would prevent cooperative growth (Figure 7b). Simulations indicated that it is the initial *E. coli* cell density rather than the initial species ratio that determines if coculture growth will be prevented through dose-dependent toxicity (Figure 7c). The inhibitory effect of high initial *E. coli* cell densities could be offset by a high initial *R. palustris* cell density enabling organic acid consumption at a rate sufficient to hamper accumulation (Figure 7c). However, simulations suggest that an initial *R. palustris* concentration of 1010 cells / ml would be required to fully offset the acidification from an initial *E. coli* cell density of 10^9^ cells / ml, mainly because NH_4_^+^cross-feeding by *R. palustris* would stimulate *E. coli* growth and thereby accelerate fermentation and organic acid accumulation. Thus, while growth-independent fermentation is a stabilizing factor at low NH_4_^+^ exchange levels, it can also serve to destabilize the mutualism at high *E. coli* densities.

**Figure 7.**
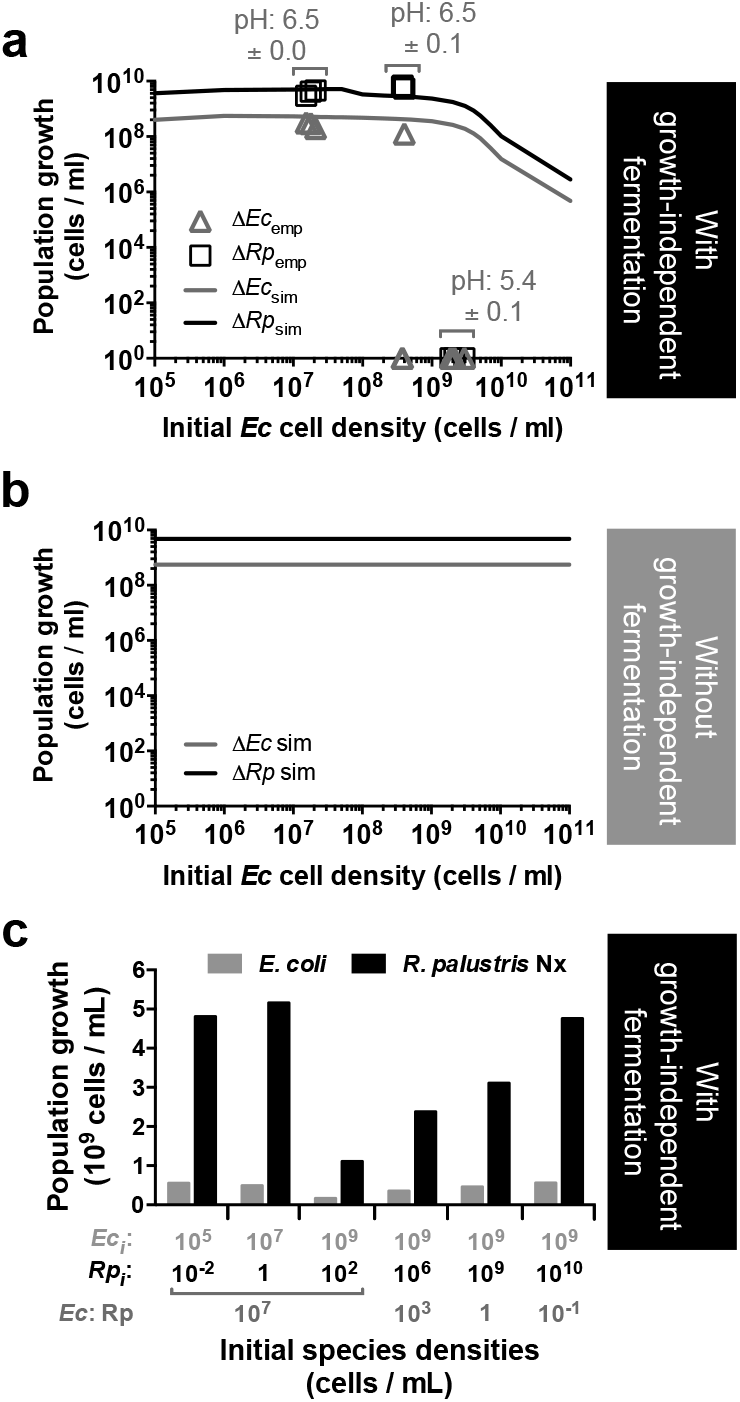
**Growth-independent fermentation prevents coculture growth at high initial *E. coli* cell densities.** (**a**, **b**) Simulated changes in *E. coli* (*Ec*sim) and *R. palustris* (*Rp*sim) cell densities (lines) in cocultures grown with 100% N_2_ from different initial *E. coli* cell densities when growth-independent fermentation is included (**a**) or omitted (**b**) from the model. The simulated initial *R. palustris* cell density was 10^7^ cells per ml. (**a**) Symbols are empirical CFUs/mL data for *E. coli* (*Ec*emp) and *R. palustris* (*Rp*emp) from cocultures with different initial E. *coli* cell densities and an average initial *R. palustris* density of 2.4 x 10^7^ ± 0.3 x 10^7^ CFU/ml. Experimental cell densities were determined 10 d after coculture inoculation. The measured coculture pH after 10 d is also shown. (**c**) Simulated *E. coli* and *R. palustris* growth when cocultures grown with 100% N_2_ are initiated from different starting cell densities. Species ratios are shown under the cell densities.

## Discussion

In this study, we demonstrated that growth-independent cross-feeding can circumstantially impede or promote mutualism. Mutualism destabilization by growth-independent metabolism depends on dose-dependent nutrient toxicity. Specifically, destabilization occurs when a cross-fed nutrient produced in a growth-independent manner accumulates sufficiently to inhibit growth of the partner species. Destabilization of a natural mutualism by growth-independent metabolism would require specific conditions. In our system, organic acid toxicity is relatively low, in part due to the buffered medium. Thus, an extremely high initial *E. coli* cell density was required before growth-independent fermentation could inhibit cooperative growth via culture acidification. However, inhibitory effects linked to growth-independent metabolism could occur at cell densities relevant to natural systems in a less well-buffered system or if the cross-fed metabolite toxicity was intrinsically high. For example, notoriously toxic compounds like cyanide (Harris and Knowles, 1983) and antibiotics (Barnhill *et al.*, 2010; Dantas *et al.*, 2008) can serve as nutrients for some bacteria as long as concentrations are maintained at low concentrations.

While the likelihood of mutualism destabilization by growth-independent metabolism is difficult to gauge, promotion of cross-feeding relationships by growth-independent metabolism is likely to be widespread. Vast areas of the Earth's biosphere are limited for key nutrients (Lever *et al.*, 2015), and it is well appreciated that nutrient limitation can promote cross-feeding in natural environments (Seth and Taga, 2014; Hom and Murray, 2014). However, it is poorly understood how established mutualisms respond to perturbations that limit cross-feeding itself. It is thought that exchange rates within obligate mutualisms must be sufficient to support sustained growth of both species in order to avoid eventual extinction (Shou *et al.*, 2007). Our results demonstrate that growth-independent cross-feeding can ease this requirement. In our system, growth-independent fermentation by *E. coli* can preserve the mutualism amid unfavorable NH_4_^+^ exchange levels by continually cross-feeding organic acids. This persistent cross-feeding stimulates *R. palustris* growth and NH_4_^+^ excretion, thereby lifting both species out of starvation. In other words, persistent growth-independent cross-feeding facilitates cooperative success over an extended range of excretion levels compared to metabolites whose excretion is dependent on growth. Given that the majority of microbes in natural environments are in a state of dormancy or low metabolic activity (Hoehler and Jørgensen, 2013; Jørgensen and Marshall, 2016; Lever *et al.*, 2015), we postulate that metabolite release is more likely to be growth-independent. As such, growth-independent metabolism could better serve to initiate and maintain mutualisms in natural environments. Separately, although growth-independent fermentation promoted growth of both partners under our study conditions, it is imaginable that mutualistic cross-feeding could purely support maintenance energy requirements in some cases, thereby promoting survival until nutrient availability improves.

Organic acids, and other fermentation products, are important metabolic intermediates in anaerobic food webs (McInerney *et al.*, 2008; Jørgensen and Marshall, 2016). Growth-independent fermentation could therefore play an important role under nutrient-limiting conditions by sustaining mutualistic relationships with acetogens, methanogens, photoheterotrophs, and anaerobically respiring microbes that rely on fermentation products for carbon and electrons. However, contributions of growth-independent metabolism to mutualisms need not be restricted to fermentation nor to natural environments. Generation of maintenance energy is likely a universally essential process. Thus, mutualistic relationships encompassing diverse lifestyles could conceivably be preserved at low metabolic rates, provided that the limiting nutrient(s) still permits the excretion of factors required to sustain partner viability. Additionally, growth-independent cross-feeding could also benefit industrial bioprocesses, which commonly use growth-limiting conditions to boost product yields. Indeed, growth-independent cross-feeding likely sustained our coculture during N_2_-limiting conditions under which the highest H_2_ yields were observed (Figures 4 and 5). Applications of microbial consortia for industrial processes is gaining interest (Sabra *et al.*, 2010) but the effects of nutrient limitation have yet to be investigated. Clearly, the role of growth-independent metabolic activities in fostering microbial cooperation deserves closer appraisal in both natural and applied systems.

## Acknowledgements

We thank David Kysela and Amelia Randich for discussions on the model name, SyFFoN. This work was supported in part by the U.S. Department of Energy, Office of Science, Office of Biological and Environmental Research, under Award Number DE-SC0008131, by the U.S. Army Research Office, grant W911NF-14-1-0411, and by the Indiana University College of Arts and Sciences.

## Supplementary Figures and Tables

**Supplementary Figure 1.**
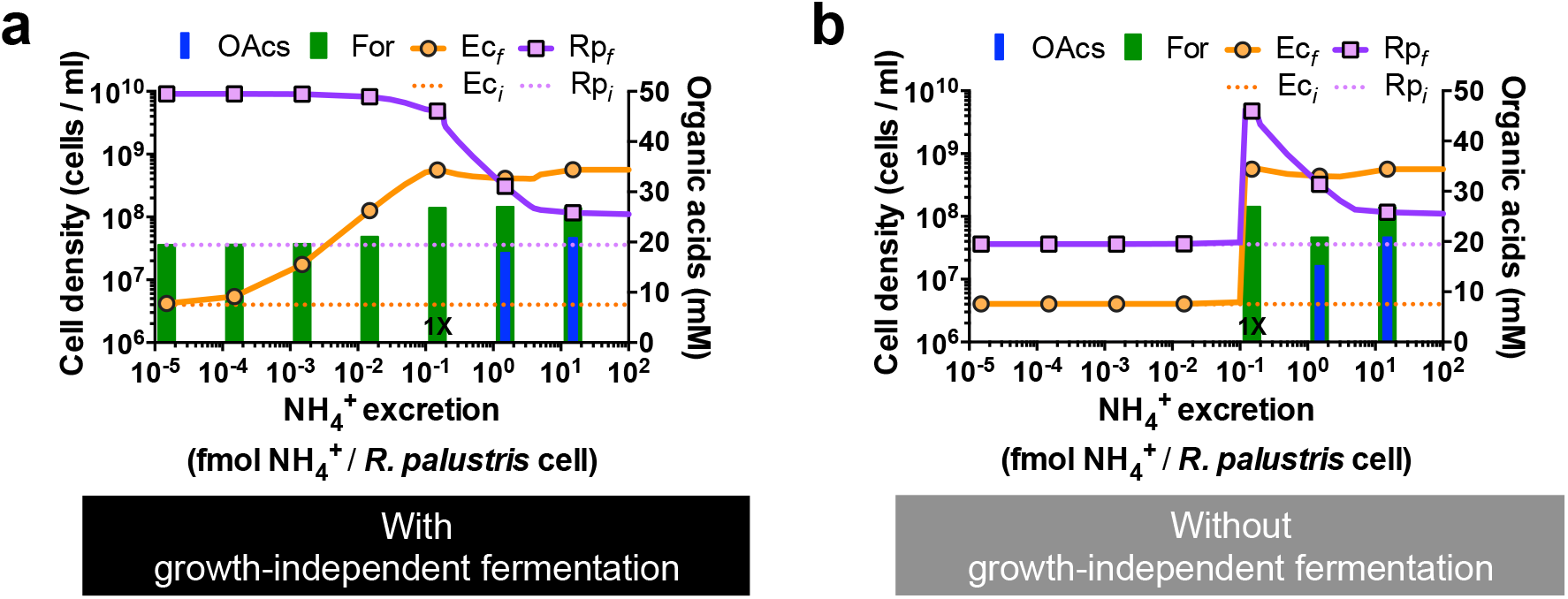
**SyFFoN_v2 predicts that coexistence at low NH 4+ cross-feeding levels requires growth-independent fermentation.** Simulated effect of the *R. palustris* NH_4_^+^ excretion level on growth and organic acid accumulation in 100% N_2_-supplied cocultures in the presence (**a**) or absence (**b**) of growth-independent fermentation. 1X is the default NH_4_^+^ excretion level (0.15 fmol NH_4_^+^ / cell) and is thought to represent that excreted by *R. palustris* Nx based on model approximation of empirical trends. OAcs, consumable organic acids (lactate, acetate, and succinate); For, formate; *Eci* and *Rpi*, initial *E. coli* (*Ec*) and *R. palustris* (*Rp*) cell densities; Ec*f* and Rp*f*, final *E. coli* and *R. palustris* cell densities. (**a**) Trends were predicted with a previous version of the model (LaSarre *et al.*, 2016).

**Supplementary Figure 2.**
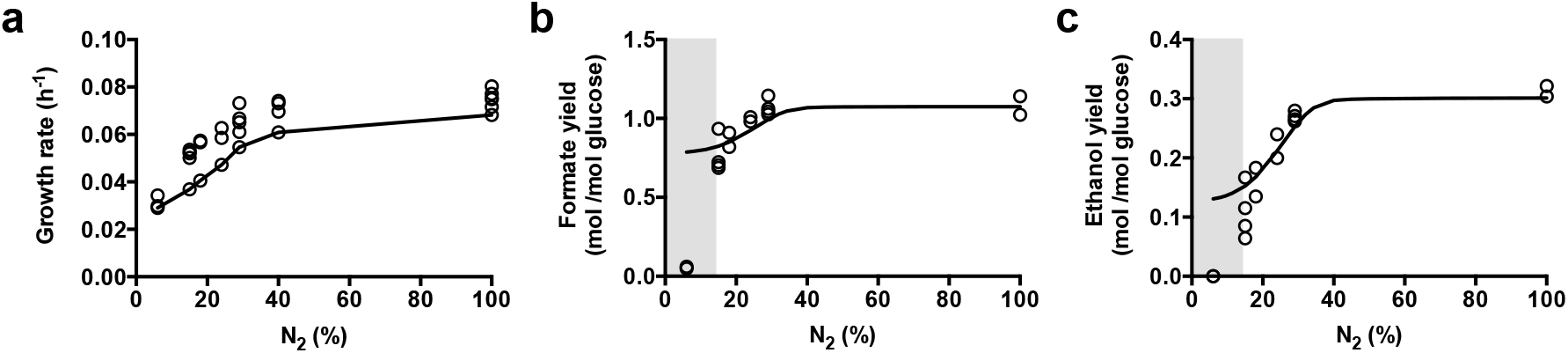
**SyFFoN_v2 predictions of growth rates (a), formate yields (b), and ethanol yields (c) at various N_2_ concentrations.** Circles indicate empirical data from shaken cocultures. Lines indicate model predictions. Shaded regions indicate low N_2_ concentrations where empirical trends do not match model predictions.

**Supplementary Figure 3.**
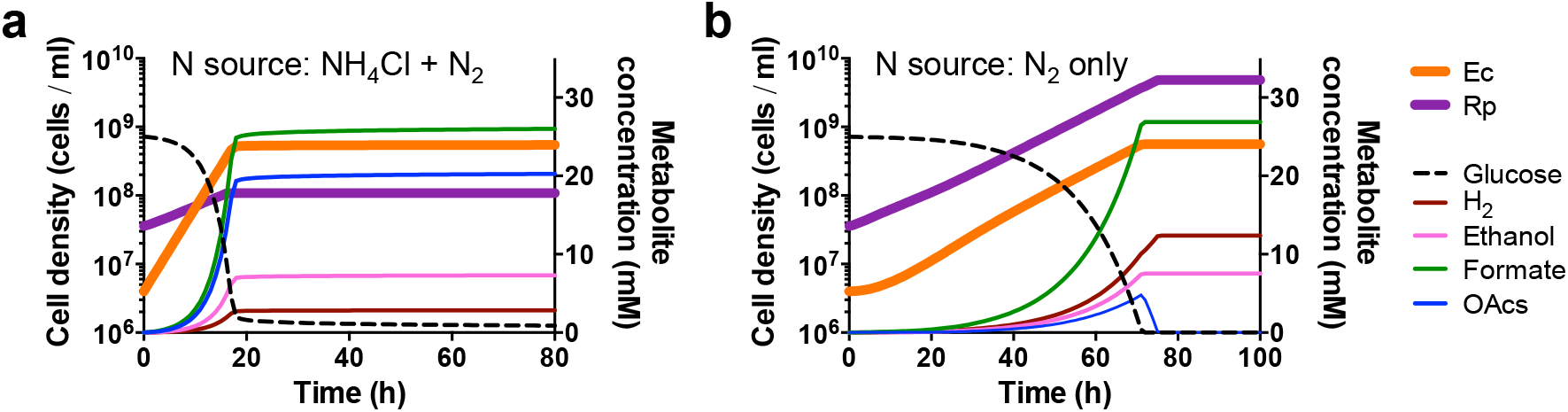
**SyFFoN_v2 simulations of batch cocultures with and without exogenously added NH_4_^+^.** (**a**, **b**) Simulated growth and metabolic profiles of cocultures supplied with NH_4_^+^ (parameter A = 15mM) (**a**) or 100% N_2_ alone (**b**). *Ec*, *E. coli*; *Rp*, *R. palustris*; OAcs, consumable organic acids (lactate, acetate, and succinate). Compare to previously published trends using an early version of the model (LaSarre *et al.*, 2016).

**Supplementary Figure 4.**
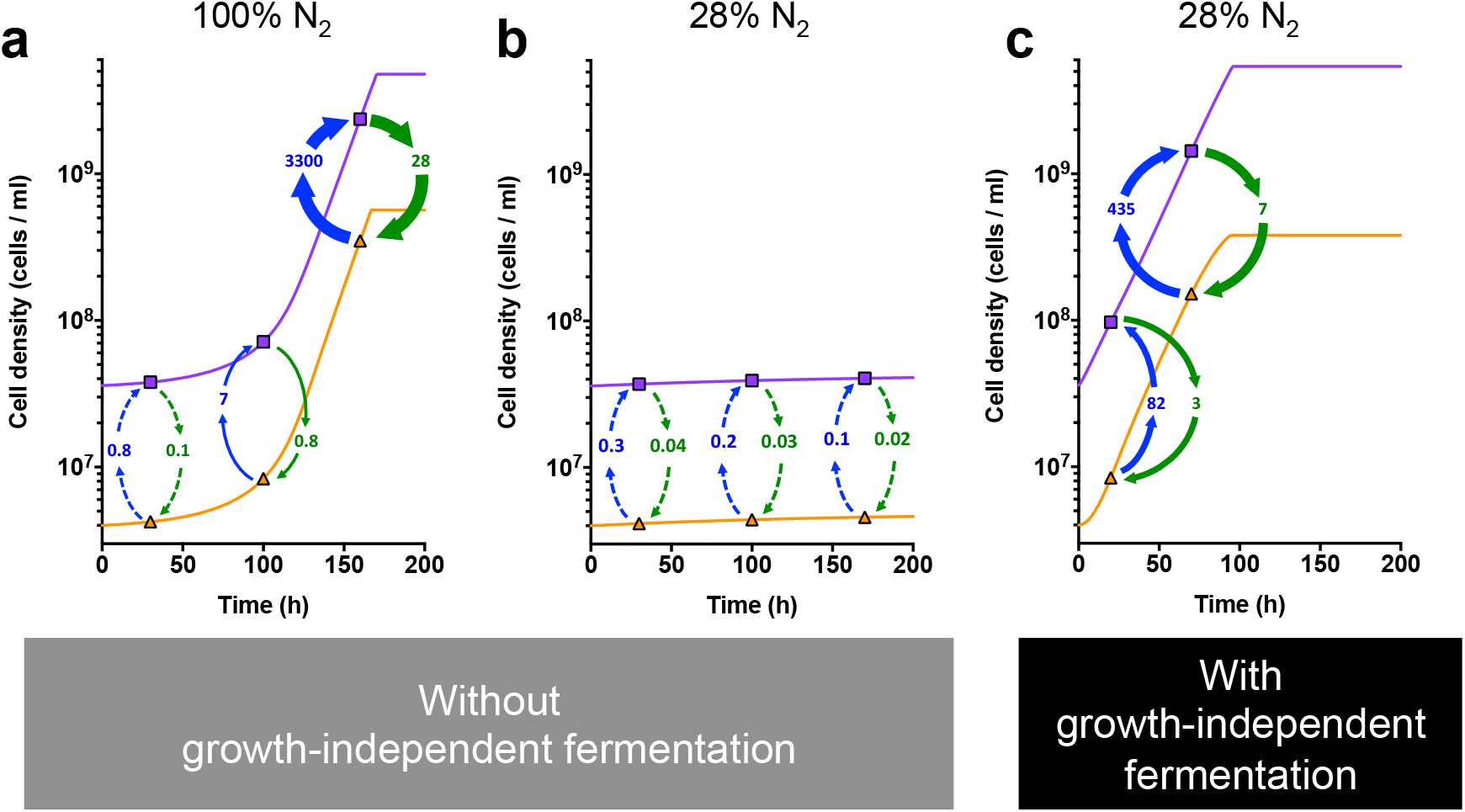
**Growth-independent fermentation is predicted to be required for coculture growth at low N_2_ concentrations.** Simulated exchange of NH_4_^+^ (green) and consumable organic acids (blue) during growth of *E. coli* (orange lines, triangles) and *R. palustris* (purple lines, squares) populations when growth-independent organic acid excretion is omitted (**a**, **b**) or included (**c**). Cross-fed NH_4_^+^ and organic acid values for the indicated time points (symbols) are the sum of those free in the medium plus those assimilated in the exchange (μM).

**Supplementary Table 1.** **Strains, plasmids and primers used in this study.**

| ***R. palustris* strains; designation in paper** |  |  |
| --- | --- | --- |
| CGA009 | Wild-type strain; spontaneous Cm<sup>R</sup> derivative of CGA001 | (Larimer *et al.*, 2004) |
| CGA4004 | CGA009 $\Delta hupS$ $\Delta rpa2750$; <u>Parent</u> | (LaSarre *et al.*, 2016) |
| CGA4005 | CGA4004 *nifA* *; <u>Nx</u> | (LaSarre *et al.*, 2016) |
| ***E. coli* strains** |  |  |
| MG1655 | Wild-type K12 obtained from the Coli Genetic Stock Center (# 7740, MG1655(Seq)), <u>WT</u> | (Blattner *et al.*, 1997) |

**Supplementary Table 2.**
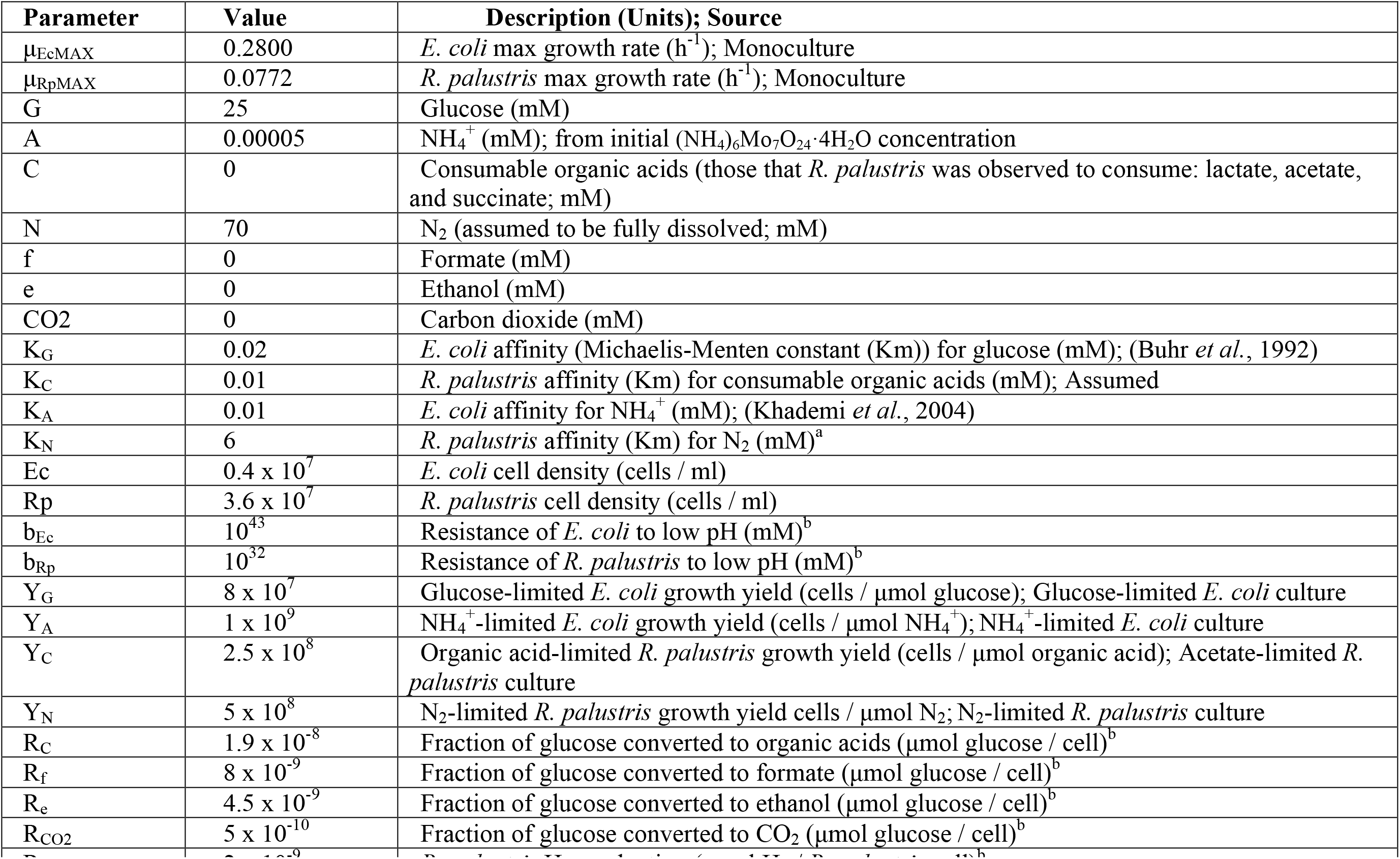
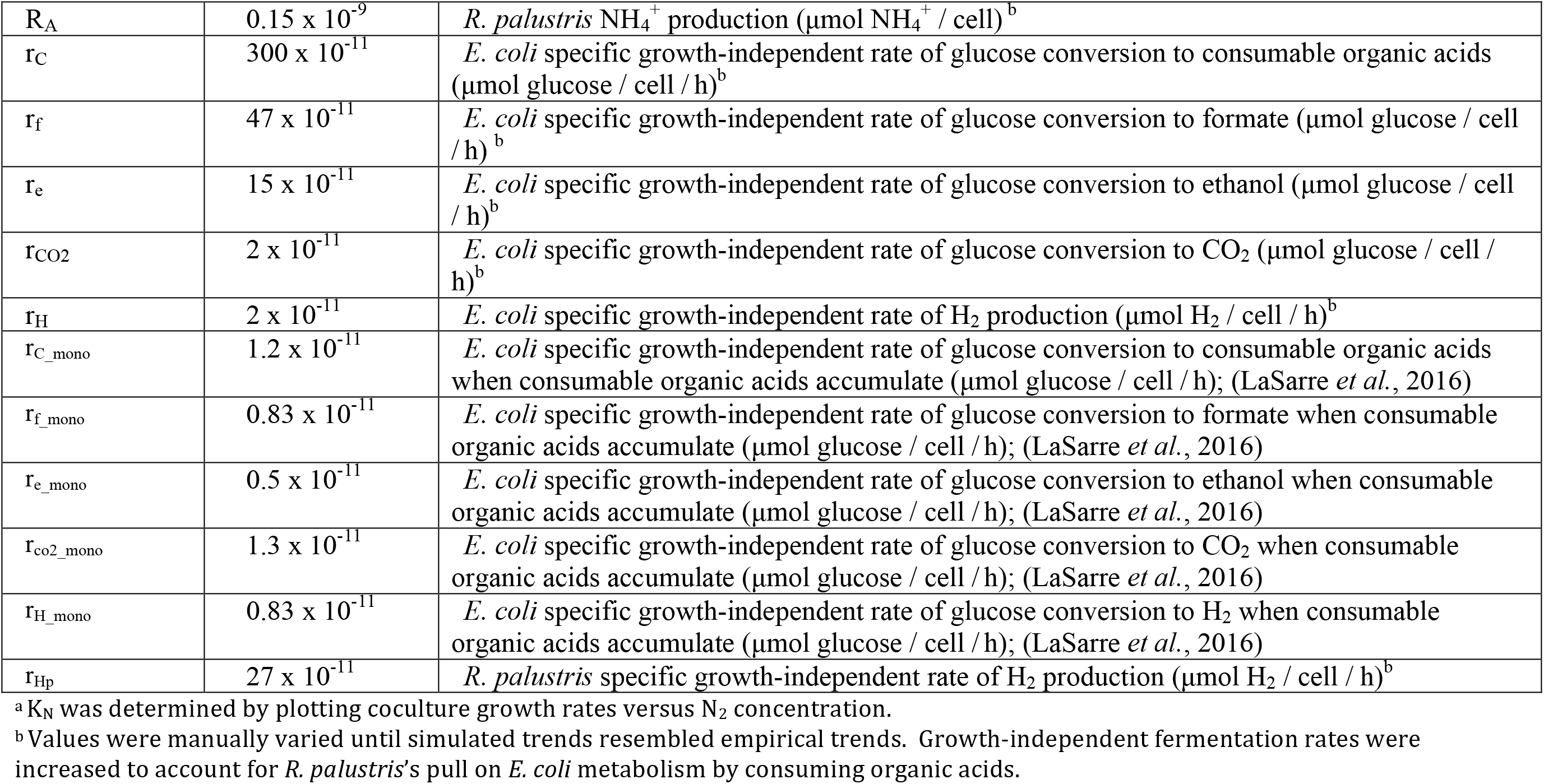
**Default parameter values used in the model unless stated otherwise**

